# Distinct Modulatory Effects on Affective Biases by Different Serotonergic Psychedelics and MDMA in Male Rats: Possible Implications for Antidepressant Effects

**DOI:** 10.64898/2026.04.20.719483

**Authors:** Justyna K Hinchcliffe, Julia Bartlett, Christopher W Thomas, Caroline T Golden, Gary Gilmour, Zuner A Bortolotto, Emma SJ Robinson

## Abstract

Affective biases are important neuropsychological mechanisms by which emotions modulate cognition, behaviour and the subjective experience of mood. Previous studies have shown that the rapid-acting antidepressant, ketamine, and serotonergic psychedelic, psilocybin, modulate affective biases in a translational rat model. Both treatments differ from conventional, delayed onset antidepressants in being able to attenuate negatively biased memories and facilitate re-learning with a more positive affective valence. Psilocybin, but not ketamine, also positively biased new experiences, an effect similar to conventional antidepressants. This study used the different affective bias test protocols, in adult male rats, to investigate the effects of acute treatment with the serotonergic psychedelics N,N-DMT, LSD and 5-MeO-DMT, and MDMA. These drugs have different pharmacology in relation to their effects on serotonin receptor subtypes and we hypothesised this may influence their modulation of affective biases. When comparing the ability to attenuate a negatively biased memory, only MDMA had specific effects although for all drugs tested, retrieval of the FG7142-induced negative affective bias was more variable and less robust statistically. LSD attenuated the negative bias at higher doses but had non-specific effects on memory retrieval. At 24hrs post treatment only N,N-DMT had a sustained effect and none of the treatments facilitated re-learning with a more positive affective valence. However, like psilocybin and conventional antidepressants, N,N-DMT positively biased new experiences. These findings suggest there are divergent affective bias modulating effects associated with different psychedelics which may be relevant to their antidepressant effects.

## Introduction

The number of patients seeking treatment for mental health conditions has risen dramatically in the last decade, exacerbated by the COVID-19 pandemic (1). Current pharmacological treatments remain limited with a significant proportion of patients failing to achieve clinically meaningful improvements in symptoms. Considering that conventional antidepressant drugs also have a delayed onset of action, there is a pressing need for better pharmacotherapies. In 2000, the NMDA receptor antagonist, ketamine, was shown to induce a rapid and sustained antidepressant effect following a single, acute low dose (2). Unlike most psychiatric drugs, ketamine’s effects endured for days after the drug had been excreted and suggest the acute treatment was able to induce longer term adaptive changes contributing to sustained effects on mood (2–4). The mechanisms underlying these sustained effects are yet to be fully elucidated but studies in animal models suggest they may be related to plasticity in brain networks associated with emotional processing (5–7).

The serotonergic psychedelics (referred to as psychedelics) are agonists or partial agonists at serotonin receptors with significant pharmacological differences within this class of drugs (8). For example, psilocin, the active metabolite of psilocybin has effects on multiple 5-HT receptor subtypes including 2A, 2B, 2C and 1A, 1B, 1D, 1E (9). N,N-DMT is relatively more potent at 5-HT1A receptors and LSD more potent at 5-HT2A receptors in humans. They also have very different pharmacokinetic profiles with the acute effects of N,N-DMT lasting less than 30min and LSD lasting for up to 8 hours (13, 14). It is not yet known the extent to which these differences in 5-HT receptor pharmacology may impact on their efficacy in mental health conditions.

In the last decade, the potential clinical benefits of serotonergic psychedelics in treating mental health disorders has seen a resurgence in interest. Clinical trials with psilocybin (15–17), DMT (18, 19) and 5-MeO-DMT (20, 21) have found positive results in treatment-resistant depression. Several countries have now permitted the medical use of psilocybin and MDMA in certain psychiatric disorders. Although not a classical psychedelic, clinical studies suggest that MDMA-assisted therapy reduces PTSD symptoms in moderate to severe PTSD (24) and reduces alcohol consumption post-detox (25). MDMA acts as an indirect serotonergic agonist, blocking the re-uptake of serotonin as well as enhancing release.

Studies in animals investigating neurotrophic effects and regional connectivity suggest that classic psychedelics promote spinogenesis, synaptogenesis, and enhance functional plasticity in cortex and hippocampus in rodent models (26–29). Investigating neurotrophic mechanisms in patients is challenging, however, imaging studies showed psilocybin induces changes in hippocampal-default mode network connectivity (30, 31). Observed changes in default mode network connectivity following psilocybin, DMT and LSD treatment may represent a neuroanatomical and mechanistic correlate of the pro-plasticity and therapeutic effects of psychedelics (30–32).

Most preclinical research relies on traditional behavioural tasks, such as the forced swim test (FST), to predict antidepressant effects, however the extent to which the FST involves mechanisms relevant to human MDD is unclear (33). The affective bias test (ABT, (34, 35)) is based on objective behavioural tasks used to quantify affective biases in human subjects (36, 37). The ABT is an associative learning task where specific cues (digging substrates) are paired with reward under different affective conditions. This generates an affective bias which can be quantified during a retrieval phase using a choice test. The ABT can be used to test if a treatment can generate an affective bias associated with new experiences or, whether treatment before the choice test, but after the induction of a biased memory, can modify retrieval. The ABT has been used to test both conventional and RAADs and the findings suggest distinct neuropsychological effects associated with affective bias modification contribute to the temporal differences in clinical effects (6, 35, 38). In our previous study investigating the effects of psilocybin in the ABT, positive biases in new learning as well as acute and sustained modulation of a negatively biased memory were observed, suggesting effects similar to both conventional antidepressants and ketamine (6).

In this study, we used the ABT and the three different testing protocols to investigate the effects of different psychedelics and MDMA on affective bias modification in male rats. We chose to test N,N-DMT, 5-MeO-DMT and LSD as these have all been trialled clinically in emotional disorders but exhibit different serotonin receptor binding profiles and duration of effects (9). We also tested MDMA given the similar mental health conditions where potential efficacy has been proposed.

## Methods

### Animals and housing

Male Lister Hooded rats (Envigo, UK) from five independent cohorts (n = 12 per cohort; details in Supplementary Table S1) were used in these experiments. At the start of training, rats were 10–11 weeks old, weighing between 300–350 g. Sample size was informed by our previous affective bias test studies and a meta-analysis (6, 39, 40). Only male subjects were used in these studies for practical reasons. Previous analysis of male and female subjects in the ABT do not suggest significant sex differences in acute pharmacological studies (39).

Animals were housed in pairs in enriched laboratory cages (55 × 35 × 21 cm) containing aspen woodchip bedding, paper bedding, a cotton rope, wood block, cardboard tube (Ø 8 cm) and a red Perspex house (30 × 17 × 10 cm). Housing was maintained under temperature-controlled conditions (21 ± 1 °C) on a 12:12 h reverse light–dark cycle (lights off at 08:00 h). Rats had *ad libitum* access to water and were maintained at ∼90% of their free-feeding weight by daily rationing of laboratory chow (∼18 g/per rat, Purina, UK) provided in the home cage at the end of each experimental day. Daily health and welfare checks were conducted by facility staff and researchers. Behavioural work was conducted during the active phase (09:00–17:00 h). All procedures complied with the UK Animals (Scientific Procedures) Act 1986 and were approved by the University of Bristol Animal Welfare and Ethical Review Body and the UK Home Office (PPL PP9516065).

### Affective Bias Test: training and testing

Before testing during week 1, rats underwent pre-training handling following the 3Hs protocol (41) and arena habituation following the protocol of Hinchcliffe & Robinson (34). In week 2, over five consecutive days, animals learned to dig in ceramic bowls filled with sawdust, with task difficulty gradually increasing. Training concluded with a novel substrate discrimination session, where rats had to learn to correctly choose and dig in the substrate associated with a food reward. ‘Correct’ choices were recorded when the reward-paired substrate was selected, ‘incorrect’ choices when the unrewarded substrate was chosen, and ‘omissions’ when the rat failed to approach within 10s. The criterion for completion was six consecutive correct trials within 20 total trials. Once successfully trained, animals undergo a reward learning assay (34) to confirm that they would develop the expected reward-induced positive bias. All animals were successfully trained and progressed to testing.

Testing weeks, each consisted of four consecutive pairing sessions generating two distinct substrate–reward associations (one under control conditions and one under treatment) (for details, see Figure 1 and Table S2). On the fifth or sixth day, a choice test was used to quantify the affective bias generated with or without drug pre-treatment (see Figure 1).

**Figure 1:**
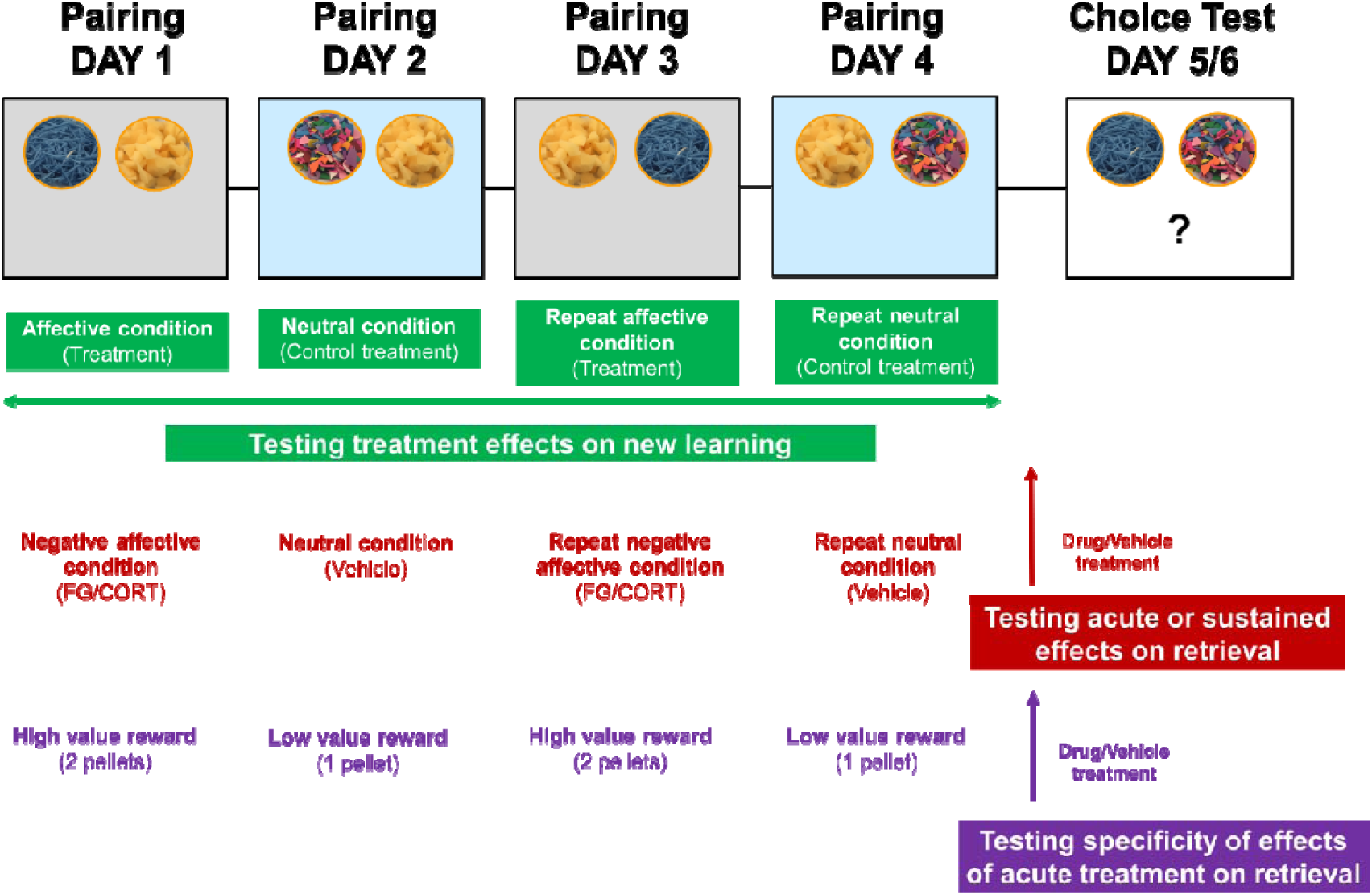
Overview of the affective bias test and reward learning assay protocols. In the affective bias test (green colour), animals are treated with either the psychedelic or vehicle prior to each of the independent substrate-reward association learning sessions with any arising affective bias quantified using a choice test 24 hours after the last pairing session. In the retrieval protocols, acute or sustained modulation of negatively biased memories (red colour) are quantified by first inducing a negative affective bias using the benzodiazepine inverse agonist FG7142 (3mg/kg) with the psychedelic or vehicle administered acutely or 24hrs before the choice test. In the reward learning assay (purple colour), animals were in the same affective state throughout training and testing but learnt to associate one of the substrate-reward cues with a higher value reward (2 reward pellets) or a low value reward (1 reward pellet). The reward-induced bias was quantified using a choice test with the RAAD or vehicle administered acutely before testing to investigate the acute effects on retrieval.

During the choice test, both previously rewarded substrates were presented simultaneously over 30 randomly reinforced trials. Choices and latency to dig were recorded. Each drug was tested as an independent experiment and a within-subject design was used with all animals receiving all treatments. The experimenter was blind to treatment, and each study used a fully counterbalanced experimental design.

### Drugs

To induce a negative affective bias in rats, the benzodiazepine inverse agonist FG7142 was used (3mg/kg, SC). The psychedelics tested were N,N-DMT (*N*,*N*-dimethyltryptamine, 0.3, 1.0 and 3.0mg/kg, IP, t=-20min and −24h), 5-MeO-DMT (5-methoxy-*N*,*N*-dimethyltryptamine, 0.03, 0.1 and 0.3mg/kg, IP, t=-40min and −24h), LSD (Lysergic acid diethylamide, 0.02, 0.03, 0.04, 0.08, 0.10, 0.16, 0.30 and 0.32 mg/kg, IP, t=-60min and −24h), MDMA (3,4-Methylenedioxymethamphetamine, 0.3, 1.0 and 3.0mg/kg, PO, t=-60min and −24h), see Table S1. Doses and route of administration were chosen based on those used clinically (18, 23, 24, 32) with the dose of LSD increased when no effects were observed. All compounds were purchased from Cambridge Bioscience. N,N-DMT was dissolved in vehicle of 5% DMSO, 10% cremophor, 85% saline and LSD in 0.9% saline. 5-Meo-DMT was initially dissolved in HCl and then 0.9% saline was added and pH adjusted. MDMA was dissolved in 1:1 strawberry milkshake (Yazoo, UK) and water. FG7142 was suspended in 0.9% saline with a drop of Tween 80 added to maintain suspension. All drugs were freshly prepared, and they were administered in a dose volume of 1.0 ml/kg. For FG7142-induced negative affective biases we selected a dose previously shown to generate a negative affective bias (6, 39). Intraperitoneal and subcutaneous injections were done by using a low-stress, non-restrained methods developed in our research group (41, 42). Prior to the start of the study, all rats were habituated to the holding positions required for IP and SC dosing and trained to voluntarily ingest the oral dosing vehicle from a 1ml syringe to facilitate subsequent oral drug dosing.

### Data analysis

Data were analysed and figures were created using GraphPad Prism 10.2.0 (GraphPad Software, USA). Percent choice bias was calculated as the number of choices made for the drug-paired substrate (affective bias test) or two pellets-paired substrate (reward learning assay) divided by the total number of trials (excluding omissions) multiplied by 100 to give a percentage. A value of 50 was then subtracted to give a score where a choice bias towards the drug-paired substrate gave a positive value and a bias towards the control-paired substrate gave a negative value. Choice bias scores and response latency scores during the choice test were analysed using a repeated measures ANOVA with treatment as the within-subject factor and Dunnett’s test for post-hoc analysis. Individual positive or negative affective biases were also analysed using a one-sample t-test against a null hypothesis mean of 0% choice bias. For each animal, mean trials to criterion, number of omissions and latency to dig during the affective bias test pairing sessions and choice test were analysed using a repeated measures ANOVA with treatment as the factor, and with post-hoc pairwise comparisons made using a two-tailed paired t-test comparison between control (vehicle/low reward:1 pellet) and treatment/manipulation (FG7142/high reward: 2 pellets) for each week. Analysis of the choice latency, omissions and trials to criterion was made to determine any non-specific effects of treatment. A Shapiro-Wilk test was used to determine a normal distribution for the % choice bias, trials to criterion, and mean latency to dig during pairing sessions and choice test. Mauchly’s sphericity test was used to validate a repeated measures ANOVA.

## Results

In the studies involving an FG7142-induced negative bias, animals that did not exhibit the expected negative bias under vehicle treatment were excluded from the whole study and any values more than 2 standard deviations from the group mean were also excluded. Three rats following the highest dose of 0.3mg/kg of 5-MeO-DMT in the acute and RLA study did not complete the choice test, therefore they were removed from the study and replaced with the group mean value to perform a full RM ANOVA. The entire treatment group receiving 3.0 mg/kg MDMA was excluded from the acute recall analysis, as 9 of the 12 animals failed to complete the choice test and showed increased number of omissions and response latencies. For reference, data from Hinchcliffe et al. (6) for psilocybin (0.3-1.0mg/kg i.p.) and ketamine (1.0mg/kg) are included in Figures 1-4.

The studies investigating acute modulation of negative affective biases are summarised in Figure 1. In this ABT design, the test drug is given after four days of learning a new negative affective bias with FG7142, on the same day as the choice test with variable treatment-to-test interval ranging from 20 – 60 minutes. In all studies in the vehicle groups, the administration of FG7142 produced a robust negative bias (N,N-DMT: p=0.0002; 5-MeO-DMT: p=0.0024; LSD (0.02-0.08mg/kg): p= 0.0013, LSD (0.16-0.32mg/kg): p=0.0001; MDMA: p=0.0001). Acute treatment with N,N-DMT, 5-MeO-DMT and low doses of LSD (0.02-0.08mg/kg) failed to attenuate the FG7142-induced negative bias. MDMA induced a significant main effect of treatment (F_₂,₂₂_ = 5.898, p=0.0001). Post hoc Dunnett tests revealed that doses of 0.3 mg/kg (p=0.0063) and 1.0 mg/kg (p=0.0374) attenuated the negative bias. The highest dose (3.0 mg/kg) was excluded from the % choice test RM ANOVA because only three animals completed the 30 trials of choice test. The MDMA treatment also affected the number of omissions (F_₃_,_₃₃_ = 5.798, p=0.0027) and choice latencies (F_₃_,_₃₃_ = 16.59, p=0.0001, Table S3). In both measures, significant differences were observed between the vehicle and highest dose (omissions: p=0.0051; latencies: p=0.0001), indicating non-specific behavioural effects and explaining why animals failed to complete the session.

The studies investigating a sustained modulation of negative affective biases are summarised in Figure 2. This test design is the same as previously described for Figure 1, except the choice test is administered 24 hours after dosing. A significant main effect of treatment was observed following N,N-DMT administration (F_₃_,_₃₃_ = 4.430, p=0.0101) 24hrs prior to the choice test. Post hoc Dunnett tests showed that doses of 1.0 mg/kg (p=0.0137) and 3.0 mg/kg (p=0.0071) attenuated the FG7142-induced negative bias but did not induce a positive bias. No main effect of treatment was observed in the 5-MeO-DMT, LSD, or MDMA studies. All vehicle groups displayed the expected FG7142-induced negative bias (5-MeO-DMT: p=0.0021; LSD: p=0.0089; MDMA: p=0.0060).

**Figure 2:**
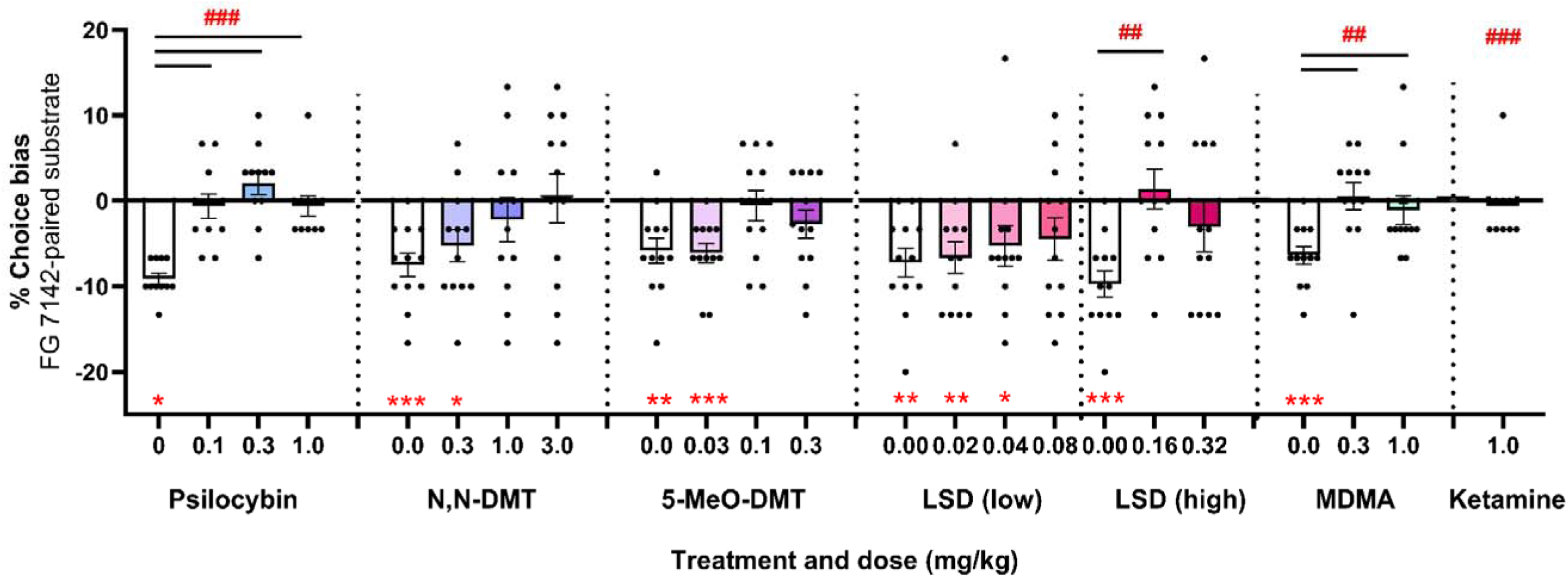
Acute treatment with MDMA but not N,N-DMT, 5-MeO-DMT or LSD attenuates negative affective biases in rats. Following induction of a negative affective bias with FG7142 male rats were injected with N,N-DMT (0.3, 1.0, 3.0mg/kg; t=-20min; n=12), 5-MeO-DMT (0.-3, 0.1, 0.3mg/kg; t=-40min; n=12), LSD (0.02, 0.04, 0.08, 0.16, 0.32mg/kg; t=-60min; n=12) or MDMA (0.3, 1.0mg/kg; t=-60min; n=12) prior to choice test. Only MDMA (0.3 and 1.0mg/kg) and high doses (0.16 and 0.32mg/kg) of LSD significantly attenuated previously learnt negatively biased memories in rats. Psilocybin and ketamine data re-drawn from Hinchcliffe et al. 2024. Data are shown as mean % choice bias ± SEM (bars) as well as individual data points (dots). Data were analyzed with one sample t-test against a null hypothesis mean of 0% choice bias (*p<0.05, **p<0.01, ***p<0.001) and pairwise comparisons were done using Dunnett’s test following main effect of treatment in RM ANOVA (##p<0.01, ###p<0.001).

The studies investigating the induction of new affective biases are summarised in Figure 3. In this ABT design, the test drug is given during the learning phase, where FG7142 was given in previous designs. Following the N,N-DMT administration in the new learning study the main effect of treatment was observed (F₄,₄₄ = 4.259, p=0.0128). Post hoc Dunnett tests indicated a difference between the vehicle group and the lowest (0.3/mg/kg: p=0.0206) and highest (3.0mg/kg: p=0.0075) doses. One-sample *t*-tests confirmed a positive bias at 0.3 mg/kg (p=0.0019), 1.0 mg/kg (p=0.0294), and 3.0 mg/kg (p=0.0002). The treatments with 5-MeO-DMT, LSD and MDMA failed to induce any biases in the new learning studies.

**Figure 3:**
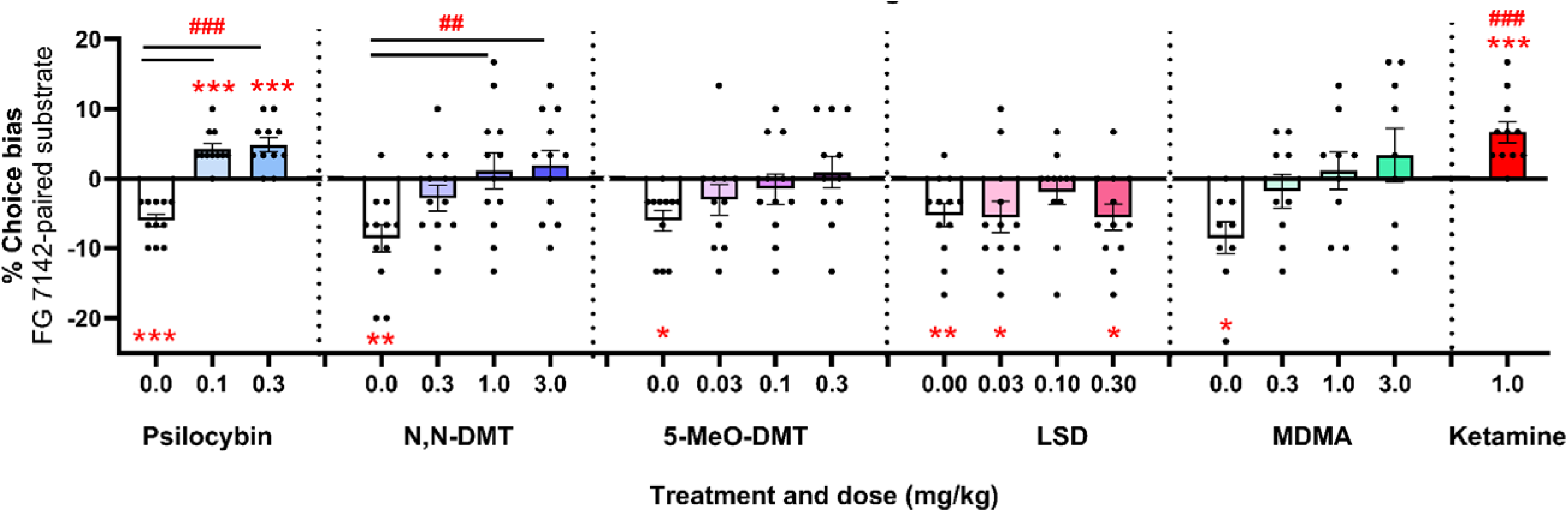
Treatment with N,N-DMT 24h prior to choice test resulted in sustained attenuation of negative affective biases in rats but no effects were seen with 5-MeO-DMT, LSD or MDMA. Following induction of a negative affective bias with FG7142 male rats were injected with N,N-DMT (0.3, 1.0, 3.0mg/kg; t=-20min; n=12), 5-MeO-DMT (0.-3, 0.1, 0.3mg/kg; t=-40min; n=11), LSD (0.02, 0.04, 0.08, 0.16, 0.32mg/kg; t=-60min; n=12) or MDMA (0.3, 1.0mg/kg; t=-60min; n=9) 24h before the choice test. Two doses of N,N-DMT (1.0mg/kg and 3.0mg/kg) significantly attenuated the negative bias. No effects were observed in other studies. Psilocybin and ketamine data re-drawn from Hinchcliffe et al. 2024. Data are shown as mean % choice bias ± SEM (bars) as well as individual data points (dots). Data were analyzed with one sample t-test against a null hypothesis mean of 0% choice bias (*p<0.05, **p<0.01, ***p<0.001) and pairwise comparisons were done using Dunnett’s test following main effect of treatment in RM ANOVA (##p<0.01, ###p<0.001).

The studies investigating the effects of the treatments on reward bias are summarised in Figure 4. In this task variant, no pharmacological treatments are used during learning, instead a positive bias is created by doubling the quantity of reward found. Testing happens after the same interval post dose as in the first experiments. No significant differences in reward-induced positive bias were observed between drug-treated animals and vehicle controls, indicating that N,N-DMT, 5-MeO-DMT, and MDMA did not impair general memory. Specifically, similar reward-induced positive biases were observed for vehicle and all N,N-DMT doses (vehicle: p=0.0004; 0.3 mg/kg: p=0.0001; 1.0 mg/kg: p=0.0015; 3.0 mg/kg: p=0.0153) and all 5-MeO-DMT doses (vehicle: p=0.0011; 0.1 mg/kg: p=0.0002; 0.3 mg/kg: p=0.0250). However, 5-MeO-DMT increased the number of omissions during the choice test (F₂,₂₂ = 4.296, p=0.0266), with post hoc Dunnett analysis showing effects at the highest dose (p=0.0409). In the MDMA study, there was no effect of treatment with similar reward-induced biases observed for all treatment groups (vehicle: p=0.0002; 0.3 mg/kg: p=0.0016; 3.0 mg/kg: p=0.0155). In contrast, LSD (0.16-0.32mg/kg) results in a significant main effect of treatment (F₂,₂₂ = 7.343, p=0.0036). Post hoc Dunnett tests revealed that doses of 0.16 mg/kg (p=0.0294) and 0.32 mg/kg (p=0.0022) impaired the reward-induced positive bias, indicating general memory disruption. LSD treatment also significantly affected choice latencies (F₂,₂₂ = 6.489, p=0.0061), with the highest dose demonstrating a significant latency increase (p=0.0038, Table S4).

**Figure 4:**
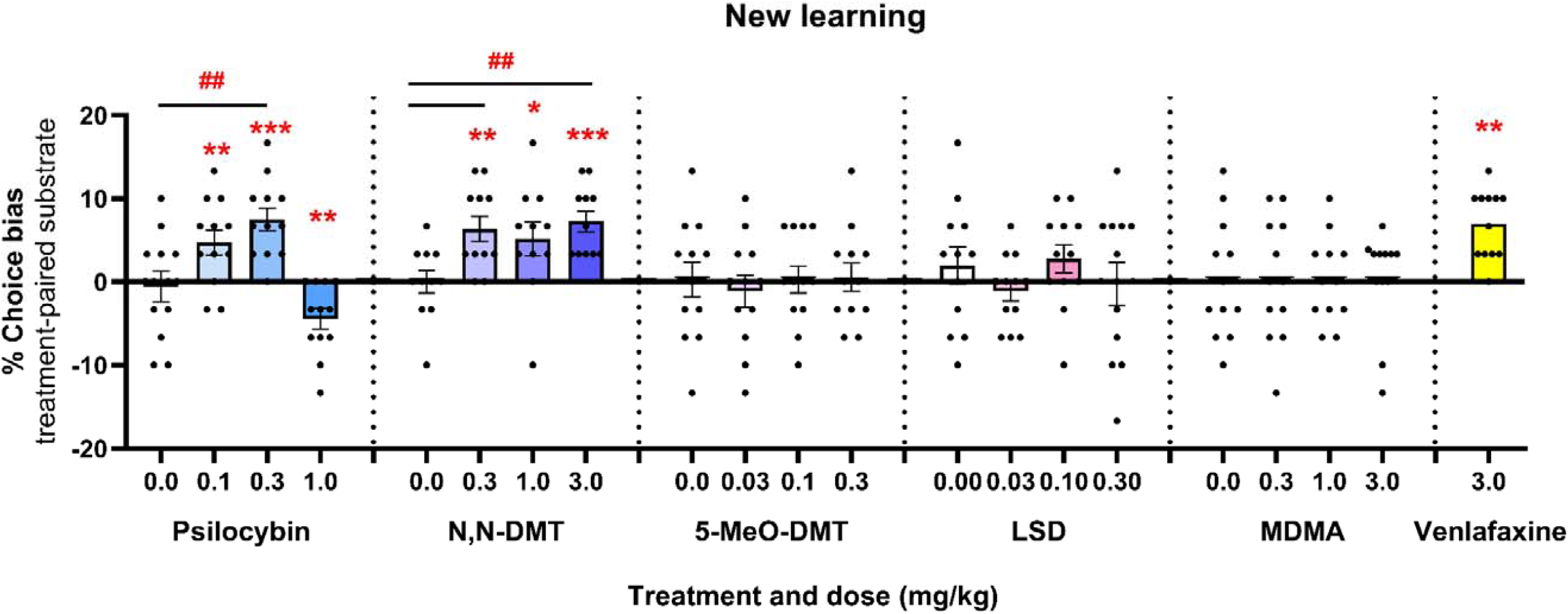
N,N-DMT induced a positive affective bias associated with new learning, but no effects were seen with 5-MeO-DMT, LSD or MDMA. Rats were acutely administered with N,N-DMT (0.3, 1.0, 3.0mg/kg; t=-20min; n=11), 5-MeO-DMT (0.03, 0.1, 0.3mg/kg; t=-40min; n=12), LSD (0.03, 0.10, 0.30mg/kg; t=-60min; n=12) or MDMA (0.3, 1.0, 3.0mg/kg; t=-60min; n=12) before learning one of the reward-paired cue associations with the choice test carried out at least 24hrs after the last treatment. A significant positive bias was only observed following treatment with all N,N-DMT doses with no effects observed for any of the other drugs tested. Psilocybin and venlafaxine data re-drawn from Hinchcliffe et al. 2024. Data are shown as mean % choice bias ± SEM (bars) as well as individual data points (dots). Data were analysed with one sample t-test against a null hypothesis mean of 0% choice bias (*p<0.05, **p<0.01, ***p<0.001) and pairwise comparisons were done using Dunnett’s test following main effect of treatment in RM ANOVA (##p<0.01).

**Figure 5:**
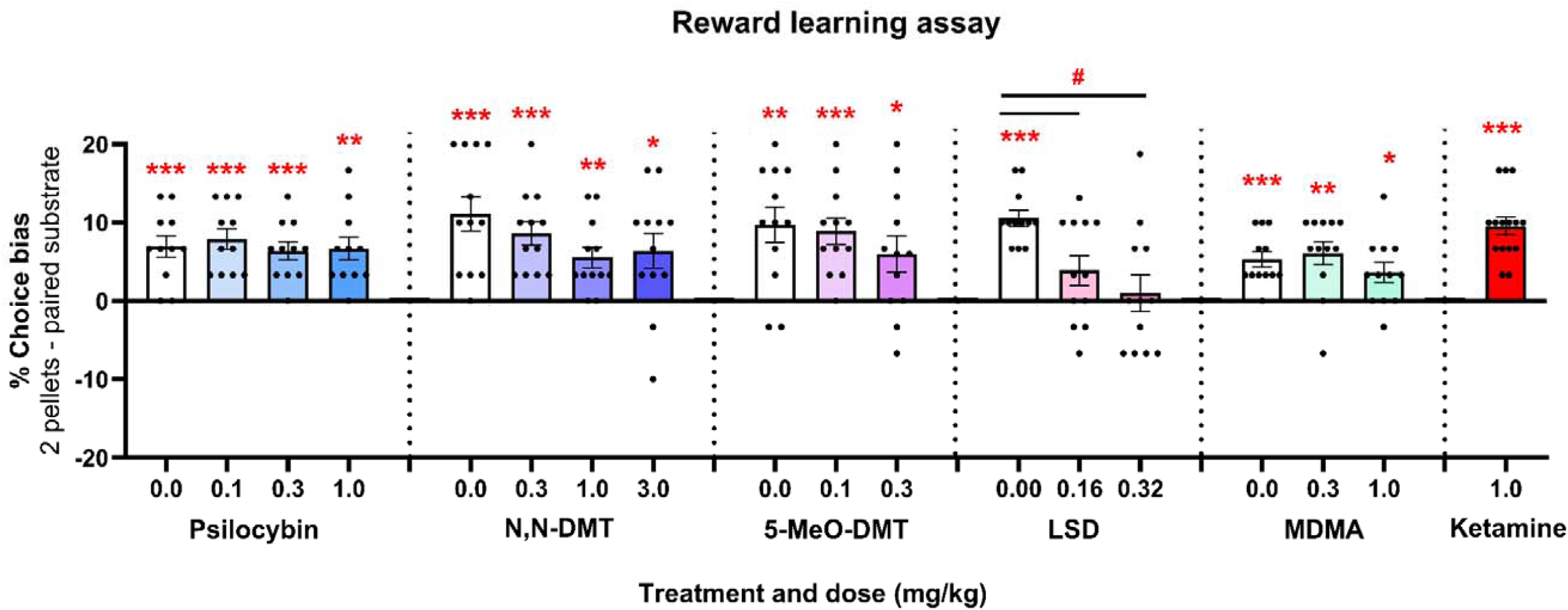
Only high doses of LSD induced general impairments resulting in non-specific effects in the reward learning assay. A reward-induced positive bias was generated by using high (two pellets) versus low (one pellet) reward outcomes associated with specific cues during the pairing sessions followed by acute administration of N,N-DMT (0.3, 1.0, 3.0mg/kg; t=-20min; n=12), 5-MeO-DMT (0.1, 0.3mg/kg; t=-40min; n=12), LSD (0.16, 0.32mg/kg; t=-60min; n=12) or MDMA (0.3, 1.0mg/kg; t=-60min; n=12) prior to the choice test. General memory impairments were observed in the study following 0.16 and 0.32mg/kg LSD treatment, indicating non-specific effects with no other effects observed. Psilocybin and ketamine data re-drawn from Hinchcliffe et al. 2024. Data are shown as mean % choice bias ± SEM (bars) as well as individual data points (dots). Data were analysed with one sample t-test against a null hypothesis mean of 0% choice bias (*p<0.05, **p<0.01, ***p<0.001) and pairwise comparisons were done using Dunnett’s test following main effect of treatment in RM ANOVA (#p<0.05).

In most of the studies there was no evidence of non-specific impairments for the psychedelic treatments tested during choice tests (Tables S3-S6) and pairing sessions (Tables S7-S10). Only changes were observed in the acute modulation studies (Table S3), during the choice test for the highest doses, 3.0mg/kg in N,N-DMT (Dunnett test, p=0.0286), 0.3mg/kg in 5-MeO-DMT (Dunnett test, p=0.0031). During pairing sessions, only treatment in the new learning study with LSD (0.10mg/kg, paired t-test, t11=2.691, p=0.0210, Table S5) resulted in slower latency to dig comparing to the vehicle group, and in the reward learning assay, treatment with 5-MeO-DMT during week 2 resulted in faster latency to dig for 2 pellets comparing to 1 pellet (paired t-test, t11=2.357, p=0.0380, Table S10).

## Discussion

These findings suggest the effects of different psychedelics on affective bias modification show significant variations which may impact on their antidepressant effects. None of the psychedelics tested exhibited the same profile of effects as psilocybin or ketamine, with only N,N-DMT demonstrating efficacy at clinically relevant doses. N,N-DMT positively biased new learning like psilocybin and conventional antidepressants, however, the effects of N,N-DMT during retrieval were limited to a sustained attenuation of negative biases. Unlike psilocybin or ketamine, N,N-DMT did not facilitate re-learning of the negatively biased memory with a more positive affective valence. Neither 5-MeO-DMT nor LSD had any effects at clinical doses, although higher doses of LSD impaired memory retrieval with non-specific effects.

The studies testing effects on retrieval of a biased memory showed higher variability and the FG7142-induced negative bias was no longer significant in the one-sample t-test. As there were no effects in the RLA, which is used to test for non-specific effects, this may reflect a specific but weaker action in terms of affective bias modulation. MDMA significantly attenuated a negative affective bias acutely with findings from the RLA suggesting these effects were specific. However, like 5-MeO-DMT and LSD, the effects at 24hrs were more variable than the vehicle group and similarly lacked a main effect of treatment. MDMA also lacked any effects on new learning. Taken together, these findings suggest the specific pharmacology of these different drugs can significantly impact their ability to modulate affective biases.

Modulation of affective biases underpins the neuropsychological hypothesis of antidepressant efficacy (36, 43). Studies in healthy volunteers and patients given acute or short-term repeated administration of conventional antidepressants suggest these drugs positively bias emotional processing and cognition (43–45). These effects enable learning with a more positive affective bias which can build over time to influence the subjective experience of mood (36, 43). Studies in human emotional processing tasks have shown that psychedelics attenuate recognition of negative facial expressions (46, 47), reduce the recognition of anger expression (48), increase positive affect (46), enhance emotional empathy, and increase prosocial behaviours (47, 49, 50). Findings from the ABT suggest the ability to modulate affective biases associated with past experiences underlies rapid alleviation of depression symptoms (6). Acute treatment-induced attenuation of a negative affective bias in the ABT has been observed with ketamine, psilocybin and scopolamine, as well as different NMDA antagonists however, only drugs which had the ability to induce sustained modulation of a biased memory at 24h are also efficacious in patients with MDD (2, 17). Like N,N-DMT, the muscarinic antagonist and putative RAAD scopolamine attenuated negative biases acutely and at 24h but did not facilitate re-learning with a more positive affective valence. In contrast, acute treatment with the NMDA antagonist PCP attenuated negative affective biases but had no effects at 24h (52). The NR2B selective NMDA antagonist CP101606 has previously been shown to have rapid and sustained antidepressant effects in patients, and in the ABT has similar effects at 24h to ketamine and psilocybin but without any acute effects (52). Given the profile of affective bias modifications seen with these RAADs, we hypothesised that the ability to facilitate re-learning with a more positive affective valence most strongly predicts enduring antidepressant effects. Based on this interpretation and the findings with conventional antidepressants and new learning, the results from the current studies suggest only N,N-DMT is likely to induce antidepressant effects via modulation of affective biases, and these may be more limited in terms of enduring effects than psilocybin. However, the acute disruption of affective biases seen with all the treatments tested may lead to short-term changes in mood. These studies were limited to investigating affective biases and other effects such as modulating anhedonia (53, 54) and cognitive flexibility (54–56) may contribute to clinical benefits.

Although all psychedelics have effects on perception and generate effects linked to their agonism or partial agonism at 5-HT2A receptors, differences in their overall receptor binding profile are well established. Considering the results from these affective bias studies, affinity for other 5-HT receptor subtypes and/or pharmacokinetic differences may be important.

Psilocybin has been shown to modulate affective biases associated with both past experiences and new learning (6). The active metabolite psilocin shows high affinity for 5-HT1 and 5-HT2 receptor subtypes and a relatively long half-life (9). In contrast, N,N-DMT has a relatively higher affinity for the 5-HT1A receptor and a much shorter duration of action. 5-MeO-DMT is similar to N,N-DMT but with a longer half-life (13, 57–59). Interpreting the LSD data is more challenging. Although the relative affinity for the 5-HT2A receptor is higher than the 5-HT1A, the rodent 5-HT2A receptor has a different amino-acid sequence associated with the ligand binding domain meaning the drug is less potent and has a shorter half-life in rodents (14, 60). Further studies using subtype selective antagonists are needed to explore the role of these different 5-HT receptors in affective bias modification.

It was somewhat surprising given the findings with conventional antidepressants that MDMA had only limited effects in the ABT. As a serotonin re-uptake inhibitor and releasing agent, it acts as an indirect serotonergic agonist, increasing synaptic concentrations of serotonin and activation of post-synaptic and pre-synaptic 5-HT receptors. Contrasting with the effects observed with serotonin specific re-uptake inhibitors, acute treatment with MDMA did not positively bias new experiences and only an acute attenuation of negative biases was observed. This suggests it can modulate affective bias circuits in the short-term but its more potent effects on serotonergic transmission does not translate to affective bias modifications suggestive of antidepressant efficacy. This may be related to the relatively high levels of serotonin released following acute treatment and widespread activation of all serotonin receptor subtypes.

Taken together, these data provide an interesting insight into differences within the class of serotonergic psychedelics in terms of affective bias modification. They also reveal how different the profile of effects of MDMA are when compared to a conventional serotonin specific re-uptake inhibitor antidepressant. The current study was limited to male rats and did not evaluate sex differences in sensitivity to the different treatments tested. We also only used healthy subjects and did not test the effects of treatment in a disease model or against different domains of impairment. These findings do not preclude the potential for these treatments to have beneficial effects on behavioural domains relevant to psychiatric disorders such as anhedonia, anxiety and fear-related behaviours related to PTSD or cognitive flexibility and behaviours relevant to substance/alcohol use disorder.

## SUPPLEMENTARY MATERIALS

**Table S1:** Summary of drug treatments in all cohorts.

| Cohort | Treatment | Dose | z | Treatment times |
| --- | --- | --- | --- | --- |
| 1-5 | FG7142<br>(benzodiazepine partial inverse agonist) | 3mg/kg | SC | 30 min prior to pairing sessions |
| 1 | N, N-DMT (*N, N*-dimethyltryptamine) | 0.3, 1.0, 3.0mg/kg | IP | 20min. (acute modulation, RLA) or 24hrs (sustained modulation) prior to choice test, 20min. prior to pairing sessions (new learning) |
| 2 | 5-MeO-DMT (5-methoxy-*N, N*-dimethyltryptamine) | 0.03, 0.1, 0.3mg/kg | IP | 40min. (acute modulation, RLA) or 24hrs (sustained modulation) prior to choice test, 40min. prior to pairing sessions (new learning) |
| 3 | LSD (Lysergic acid diethylamide) | 0.02, 0.03, 0.04, 0.08, 0.10, 0.16, 0.30, 0.32 mg/kg | IP | 60min. (acute modulation, RLA) or 24hrs (sustained modulation) prior to choice test, 60min. prior to pairing sessions (new learning) |
| 2, 4, 5 | MDMA (3,4-Methylenedioxymethamphetamine) | 0.3, 1.0, 3.0mg/kg | PO | 60min. (acute modulation, RLA) or 24hrs (sustained modulation) prior to choice test, 60min. prior to pairing sessions (new learning) |

**Table S2:** List of the example substrates used in the experiments in both cohorts.

|  | Substrate 'A' | Substrate 'B' | Substrate 'C' |
| --- | --- | --- | --- |
| **Test 1** | fur | polyester | pompoms |
| **Test 2** | cellulose sponge | corrugated paper | perlite |
| **Test 3** | purple ribbon | green raffia ribbon | sparkly pompoms |
| **Test 4** | brown pet bedding | cork | hessian sack |
| **Test 5** | cotton wool balls | stringy cloth | hairbands |
| **Test 6** | organza | silk | shredded paper |
| **Test 7** | bin liner | plastic scourer | straws |
| **Test 8** | cotton mix | leather | balloons |
| **Test 9** | chubby wool | shoe laces | velcro |
| **Test 10** | brown partition paper | dishcloth squares | polyester lining |
| **Test 11** | aspen | cypress | coloured matchsticks |
| **Test 12** | Christmas ribbon | umbrella | tights |
| **Test 13** | towel | canvas | pipe cleaners |
| **Test 14** | newspaper | paper pet bedding | confetti |
| **Test 15** | suede | chenille strands | yellow fleece |
| **Test 16** | poster squares | polystyrene | sequins |
| **Test 17** | crepe paper squares | scarf yarn | sparkling fibre |
| **Test 18** | denim | rucksack strap | foam pad |
| **Test 19** | dusters | tissue paper balls | yellow bath sponge |
| **Test 20** | black satin | cardboard | rope |

**Table S3:**
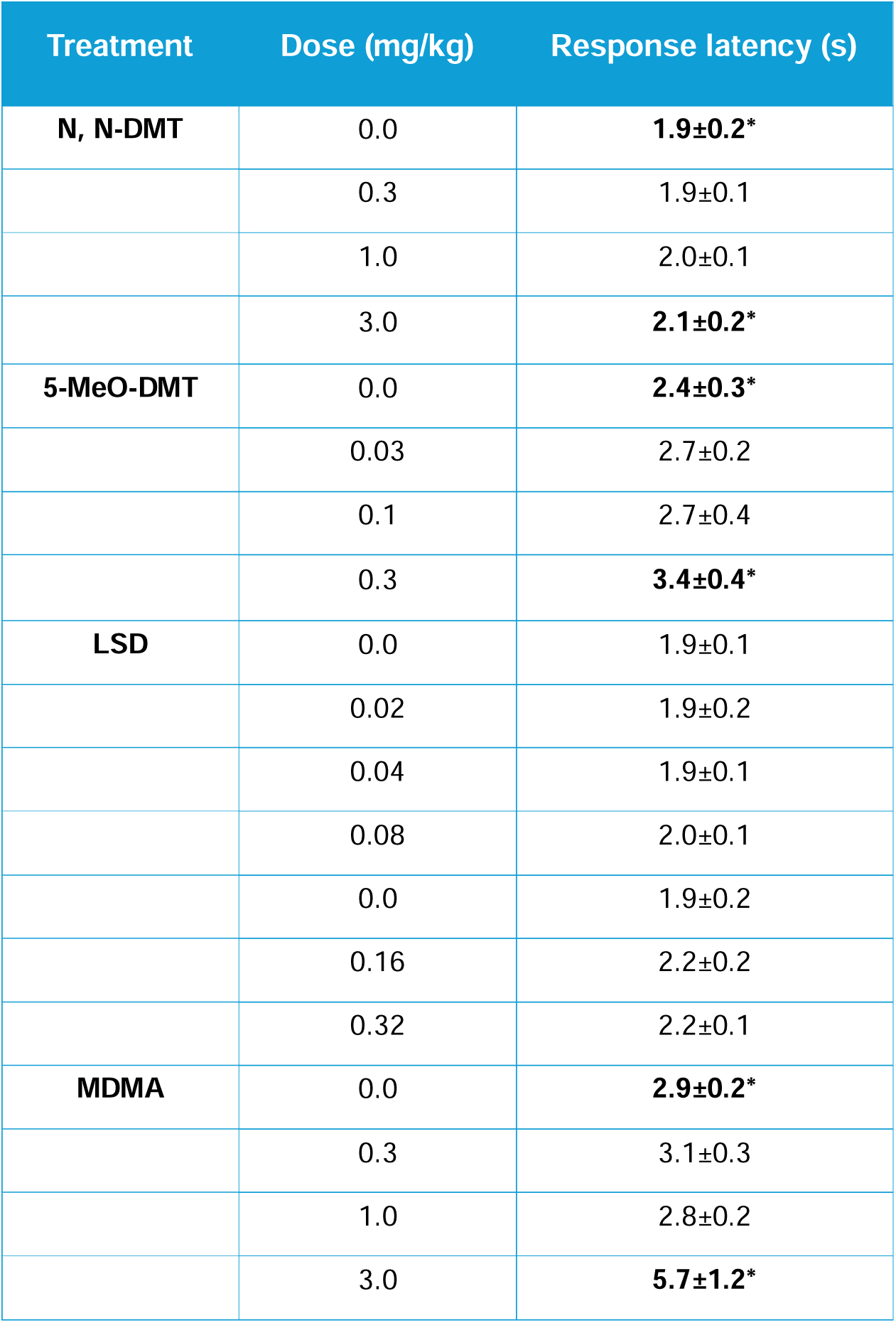
Choice bias data: response latency to make choice in the acute modulation studies. Data shown as mean (n=12 animals/group) ± SEM of an individual latencies during 30 trials of the choice test. Following treatment with N,N-DMT, 5-MeO-DMT and MDMA the latencies during choice test were altered. In all studies, the highest doses, 3.0mg/kg in N,N-DMT study (Dunnett test, p=0.0286), 0.3mg/kg in 5-MeO-DMT study (Dunnett test, p=0.0031) and 3.0mg/kg in MDMA study significantly increased latencies (Dunnett test, p=0.0027). No significant effects were observed in the LSD study.

**Table S4:**
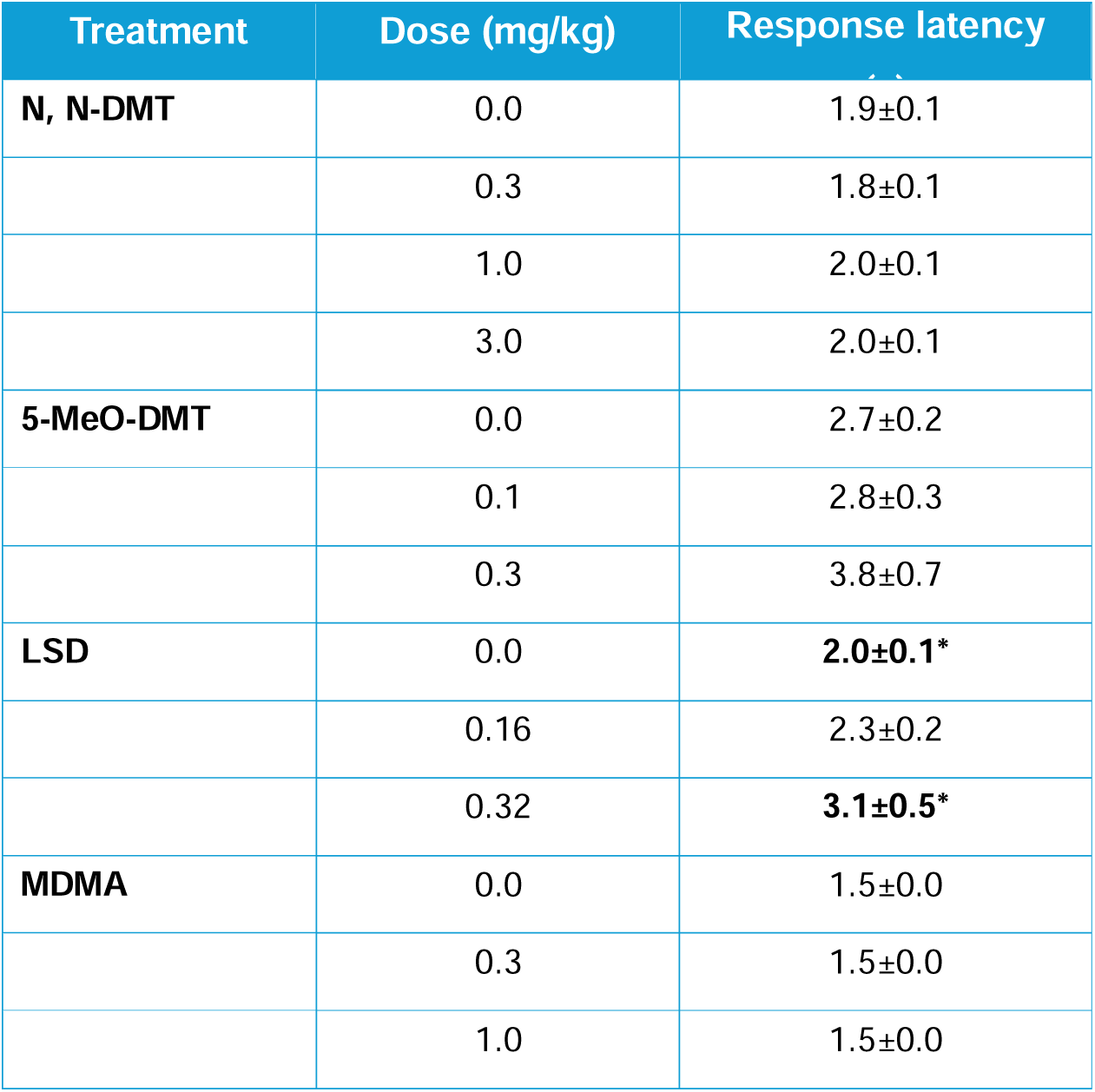
Choice bias data: response latency to make choice in the reward learning assay studies. Data shown as mean (n=12 animals/group) ± SEM of an individual latencies during 30 trials of the choice test. No significant difference in latency to make choice was observed in studies following treatment with vehicle or any of the drugs: N,N-DMT (0.3-3.0mg/kg), 5-MeO-DMT (0.1-0.3mg/kg) and MDMA (0.3-1.0mg/kg). The only significant difference was observed in the LSD study, following the highest dose 0.32mg/kg the animals were slower comparing to the vehicle group (p=0.0038).

**Table S5:** Choice bias data: response latency to make choice in the sustained modulation studies. Data shown as mean (n=9-12 animals/group) ± SEM of an individual latencies during 30 trials of the choice test. No significant difference in latency to make choice was observed following any of the treatments.

| Treatment | Dose (mg/kg) | Response latency (s) |
| --- | --- | --- |
| **N, N-DMT** | 0.0 | 1.9±0.1 |
|  | 0.3 | 1.9±0.1 |
|  | 1.0 | 1.9±0.1 |
|  | 3.0 | 2.0±0.1 |
| **5-MeO-DMT** | 0.0 | 2.8±0.2 |
|  | 0.03 | 2.8±0.2 |
|  | 0.1 | 2.7±0.3 |
|  | 0.3 | 3.3±0.6 |
| **LSD** | 0.0 | 2.0±0.1 |
|  | 0.03 | 2.1±0.1 |
|  | 0.10 | 2.2±0.1 |
|  | 0.30 | 2.1±0.1 |
| **MDMA** | 0.0 | 2.0±0.2 |
|  | 0.3 | 2.0±0.2 |
|  | 1.0 | 2.1±0.2 |
|  | 3.0 | 2.1±0.2 |

**Table S6:**
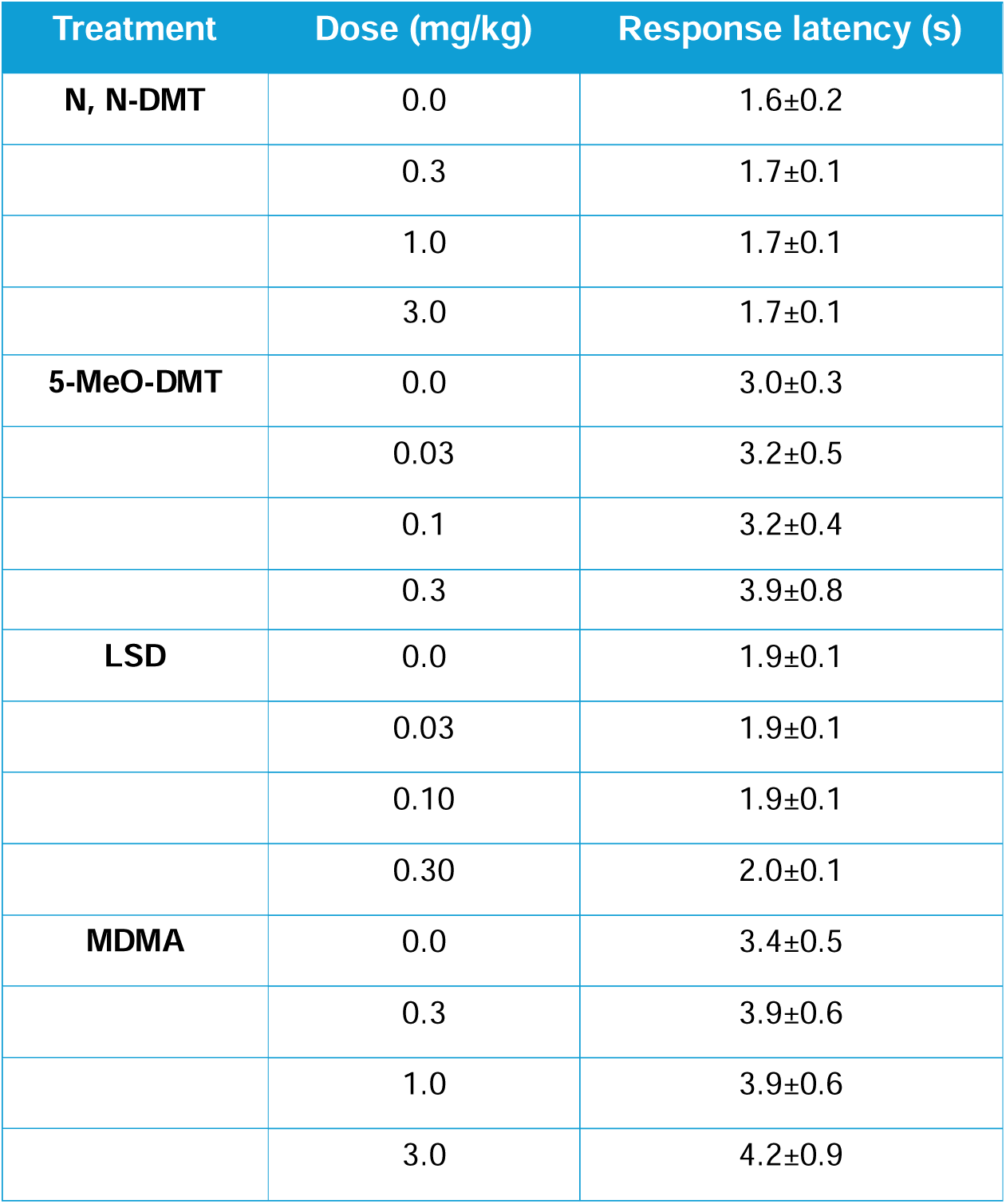
Choice bias data: response latency to make choice in the new learning studies. Data shown as mean (n=11-12 animals/group) ± SEM of an individual latencies during 30 trials of the choice test. No significant difference in latency to make choice was observed following any of the treatments.

**Table S7:**
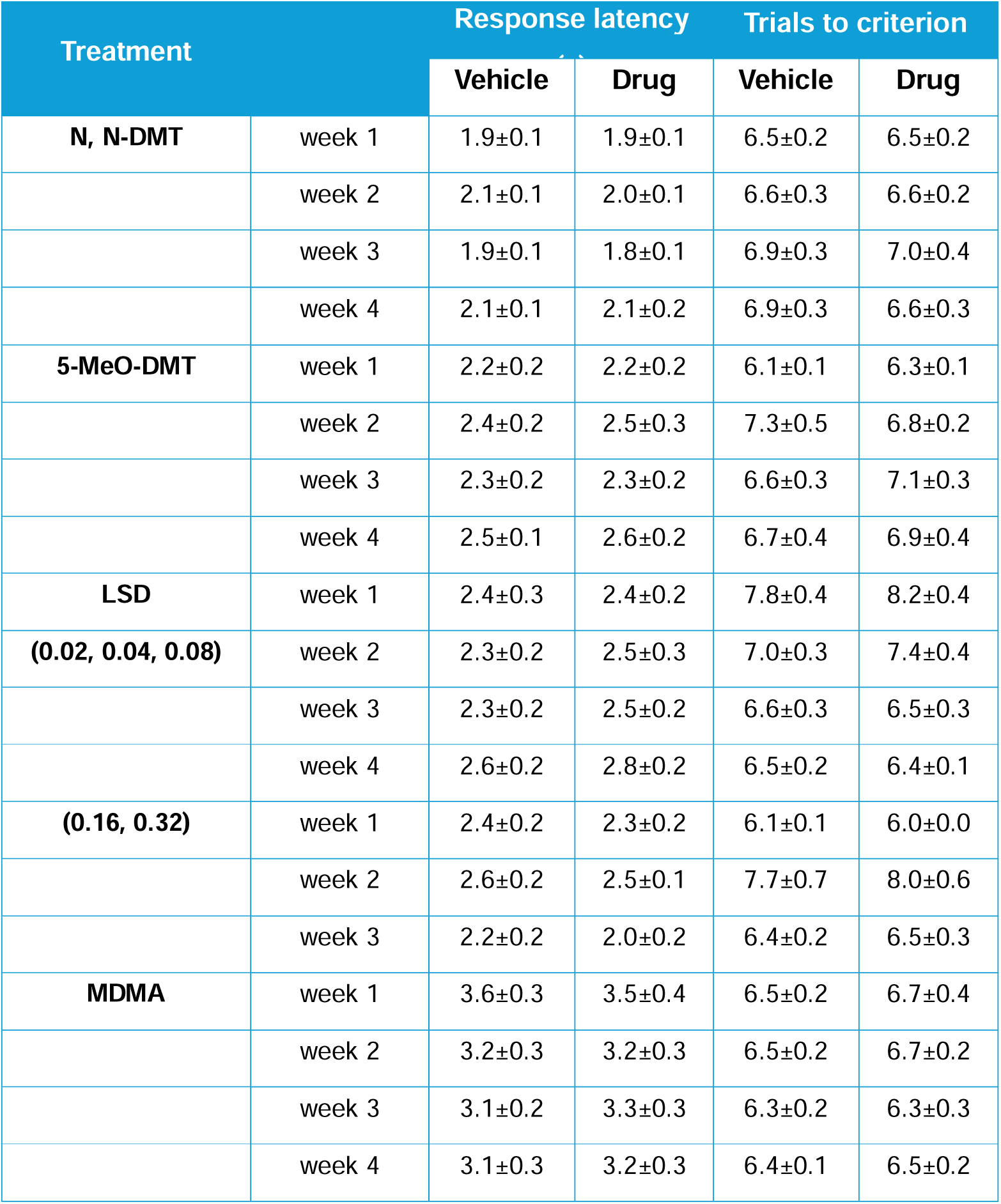
Pairing sessions data: number of trials to criterion and latency to dig in the acute modulation studies. Data shown as mean (n=12 animals/group) ± SEM averaged from the two pairing sessions for each substrate-reward association (vehicle or drug). There were no significant effects during pairing sessions, either on response latency to dig or number of trials to criterion in any of the treatment groups.

**Table S8:** Pairing sessions data: number of trials to criterion and latency to dig in the sustained modulation studies. Data shown as mean (n=9-12 animals/group) ± SEM averaged from the two pairing sessions for each substrate-reward association (vehicle or drug). There were no significant effects during pairing sessions, either on response latency to dig or number of trials to criterion in any of the treatment groups.

| Treatment |  | Response latency (s) |  | Trials to criterion |  |
| --- | --- | --- | --- | --- | --- |
|  |  | Vehicle | Drug | Vehicle | Drug |
| **N, N-DMT** | week 1 | 2.2±0.1 | 2.2±0.2 | 6.6±0.1 | 7.2±0.4 |
|  | week 2 | 1.7±0.1 | 1.7±0.1 | 6.5±0.2 | 6.5±0.3 |
|  | week 3 | 1.8±0.1 | 1.8±0.1 | 6.3±0.2 | 6.3±0.2 |
|  | week 4 | 1.8±0.1 | 1.7±0.1 | 6.3±0.1 | 6.4±0.2 |
| **5-MeO-DMT** | week 1 | 2.7±0.3 | 3.0±0.3 | 8.0±0.5 | 7.1±0.4 |
|  | week 2 | 2.8±0.4 | 2.9±0.2 | 6.5±0.2 | 6.3±0.2 |
|  | week 3 | 2.7±0.2 | 2.5±0.2 | 7.0±0.4 | 7.3±0.4 |
|  | week 4 | 2.2±0.2 | 2.2±0.2 | 6.7±0.3 | 6.7±0.3 |
| **LSD** | week 1 | 2.5±0.3 | 2.5±0.1 | 7.0±0.4 | 6.9±0.3 |
|  | week 2 | 2.7±0.6 | 2.1±0.1 | 6.9±0.3 | 6.8±0.2 |
|  | week 3 | 2.0±0.1 | 2.0±0.1 | 6.3±0.2 | 6.3±0.1 |
|  | week 4 | 1.9±0.1 | 2.1±0.1 | 6.7±0.2 | 6.8±0.3 |
| **MDMA** | week 1 | 2.1±0.1 | 2.0±0.1 | 8.5±0.7 | 7.7±0.4 |
|  | week 2 | 2.1±0.1 | 2.0±0.1 | 6.4±0.2 | 6.3±0.3 |
|  | week 3 | 2.1±0.1 | 2.0±0.1 | 6.5±0.2 | 6.5±0.3 |
|  | week 4 | 2.4±0.1 | 2.5±0.2 | 6.7±0.2 | 7.1±0.4 |

**Table S9:** Pairing sessions data: number of trials to criterion and latency to dig in the new learning studies. Data shown as mean (n=11-12 animals/group) ± SEM averaged from the two pairing sessions for each substrate-reward association (vehicle or drug). There were no significant effects during pairing sessions, either on response latency to dig or number of trials to criterion following treatment with vehicle or N,N-DMT (0.3-3.0mg/kg), 5-MeO-DMT (0.03-0.3mg/kg) and MDMA (0.3-3.0mg/kg). Only treatment with LSD (0.10mg/kg, paired t-test, t11=2.691, p=0.0210) resulted in slower latency to dig comparing to the vehicle group.

| Treatment |  | Response latency (s) |  | Trials to criterion |  |
| --- | --- | --- | --- | --- | --- |
|  |  | Vehicle | Drug | Vehicle | Drug |
| **N, N-DMT** | 0.0 | 1.7±0.1 | 1.8±0.1 | 6.3±0.2 | 6.4±0.2 |
|  | 0.3 | 1.7±0.1 | 1.9±0.2 | 6.6±0.2 | 6.8±0.4 |
|  | 1.0 | 1.9±0.1 | 1.6±0.1 | 6.8±0.4 | 6.1±0.1 |
|  | 3.0 | 1.7±0.1 | 1.7±0.1 | 6.2±0.1 | 6.4±0.2 |
| **5-MeO-DMT** | 0.0 | 2.8±0.2 | 2.7±0.4 | 6.7±0.2 | 6.6±0.3 |
|  | 0.03 | 2.5±0.2 | 2.5±0.2 | 6.6±0.2 | 7.0±0.3 |
|  | 0.1 | 2.8±0.5 | 2.7±0.4 | 6.2±0.1 | 6.5±0.2 |
|  | 0.3 | 3.0±0.4 | 4.5±1.0 | 7.0±0.3 | 6.5±0.2 |
| **LSD** | 0.0 | 1.8±0.1 | 2.1±0.1 | 6.6±0.3 | 6.5±0.2 |
|  | 0.03 | 2.2±0.1 | 2.1±0.1 | 6.6±0.3 | 7.0±0.4 |
|  | 0.10 | **1.8±0.1\*** | **2.2±0.2\*** | 6.3±0.2 | 7.0±0.5 |
|  | 0.30 | 2.2±0.1 | 2.5±0.2 | 6.6±0.2 | 6.5±0.2 |
| **MDMA** | 0.0 | 3.4±0.6 | 3.6±0.9 | 6.5±0.2 | 6.3±0.2 |
|  | 0.3 | 3.1±0.3 | 2.9±0.3 | 6.7±0.3 | 6.2±0.1 |
|  | 1.0 | 3.3±0.4 | 3.4±0.4 | 6.6±0.3 | 6.4±0.2 |
|  | 3.0 | 3.9±1.0 | 5.1±1.0 | 6.5±0.2 | 6.3±0.2 |

**Table S10:**
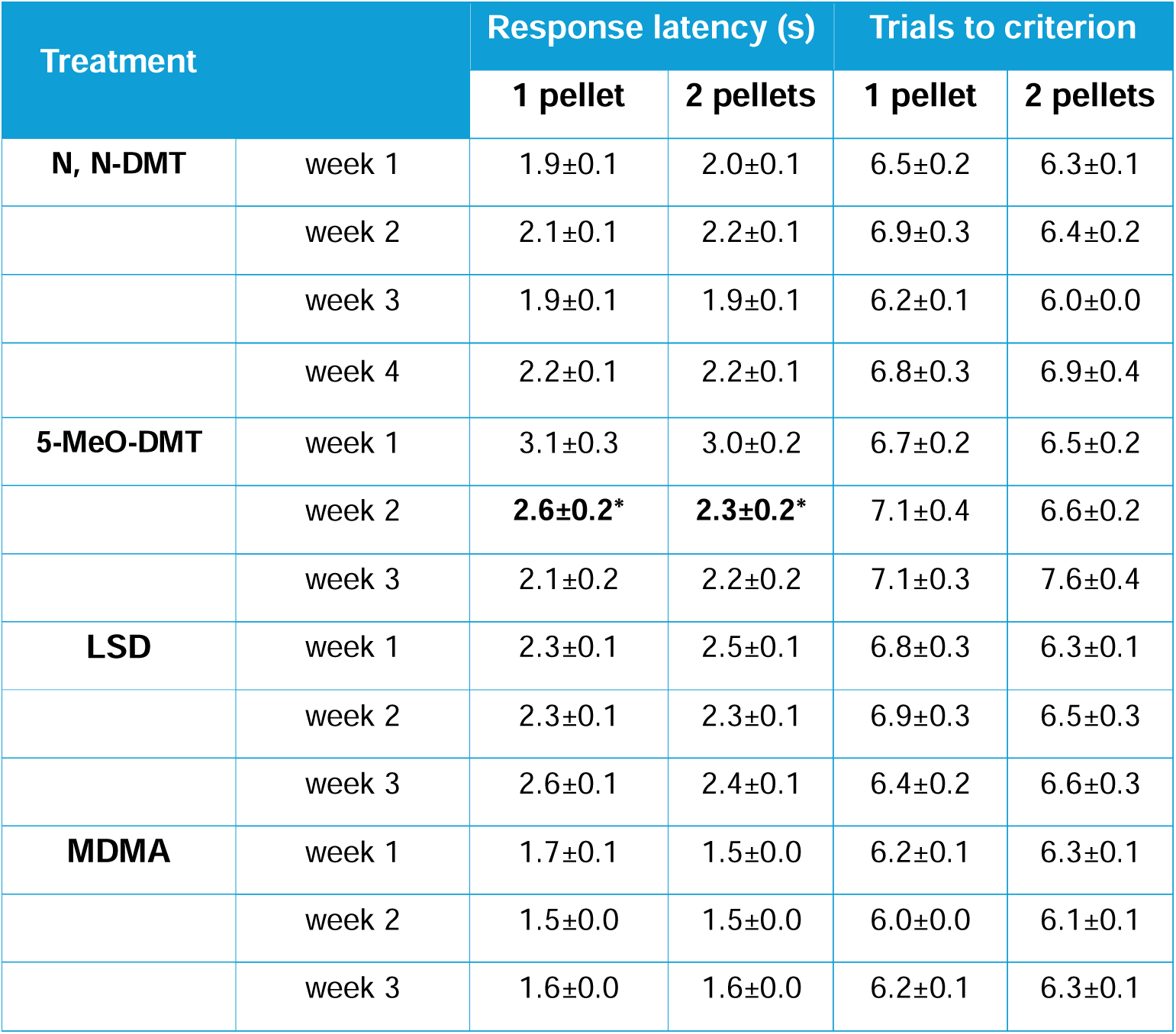
Pairing sessions data: number of trials to criterion and latency to dig in the reward learning assay studies. Data shown as mean (n=12 animals/group) ± SEM averaged from the two pairing sessions for each substrate-reward association (1 pellet or 2 pellets). There were no significant effects during pairing sessions, either on response latency to dig or number of trials to criterion following treatment with vehicle or N,N-DMT (0.3-3.0mg/kg), LSD (0.03-0.30mg/kg) and MDMA (0.3-3.0mg/kg). Only treatment with 5-MeO-DMT during week 2 resulted in faster latency to dig for 2 pellets comparing to 1 pellet (paired t-test, t11=2.357, p=0.0380).

## References

1. Adult Psychiatric Morbidity Survey: Survey of Mental Health and Wellbeing, England, 2023/4 2025 [Available from: https://www.bma.org.uk/advice-and-support/nhs-delivery-and-workforce/pressures/mental-health-pressures-data-analysis.

2. Berman RM, Cappiello A, Anand A, Oren DA, Heninger GR, Charney D, et al. Antidepressant effects of ketamine in depressed patients. Biological psychiatry. 2000;47(4):351–4.

3. Kim JW, Suzuki K, Kavalali ET, Monteggia LM. Bridging rapid and sustained antidepressant effects of ketamine. Trends Mol Med. 2023;29(5):364–75.

4. Zarate CA, Jr., Singh JB, Carlson PJ, Brutsche NE, Ameli R, Luckenbaugh DA, et al. A randomized trial of an N-methyl-D-aspartate antagonist in treatment-resistant major depression. Arch Gen Psychiatry. 2006;63(8):856–64.

5. Autry AE, Adachi M, Nosyreva E, Na ES, Los MF, Cheng PF, et al. NMDA receptor blockade at rest triggers rapid behavioural antidepressant responses. Nature. 2011;475(7354):91–5.

6. Hinchcliffe JK, Stuart SA, Wood CM, Bartlett JM, Kamenish K, Arban R, et al. Rapid-acting antidepressant drugs modulate affective bias in rats. Science Translational Medicine. 2024;16(729):eadi2403.

7. Wohleb ES, Franklin T, Iwata M, Duman RS. Integrating neuroimmune systems in the neurobiology of depression. Nature reviews Neuroscience. 2016;17(8):497–511.

8. Vollenweider FX, Preller KH. Psychedelic drugs: neurobiology and potential for treatment of psychiatric disorders. Nature Reviews Neuroscience. 2020;21(11):611–24.

9. Erkizia-Santamaría I, Alles-Pascual R, Horrillo I, Meana JJ, Ortega JE. Serotonin 5-HT(2A), 5-HT(2c) and 5-HT(1A) receptor involvement in the acute effects of psilocybin in mice. In vitro pharmacological profile and modulation of thermoregulation and head-twich response. Biomedicine & pharmacotherapy = Biomedecine & pharmacotherapie. 2022;154:113612.

10. Hayes C. A Toad Less Traveled: Should 5-MeO-DMT Have a Role in Treating Depression? Psychedelic Medicine.0(0):null.

11. Hong-Wu S, Xi-Ling J, Jerrold CW, Ai-Ming Y. Psychedelic 5-Methoxy-N,N-Dimethyltryptamine: Metabolism, Pharmacokinetics, Drug Interactions, and Pharmacological Actions. Current Drug Metabolism. 2010;11(8):659–66.

12. Rickli A, Moning OD, Hoener MC, Liechti ME. Receptor interaction profiles of novel psychoactive tryptamines compared with classic hallucinogens. European Neuropsychopharmacology. 2016;26(8):1327–37.

13. Carbonaro TM, Gatch MB. Neuropharmacology of N,N-dimethyltryptamine. Brain research bulletin. 2016;126(Pt 1):74–88.

14. Passie T, Halpern JH, Stichtenoth DO, Emrich HM, Hintzen A. The Pharmacology of Lysergic Acid Diethylamide: A Review. CNS Neuroscience & Therapeutics. 2008;14(4):295–314.

15. Aaronson ST, van der Vaart A, Miller T, LaPratt J, Swartz K, Shoultz A, et al. Single-Dose Psilocybin for Depression With Severe Treatment Resistance: An Open-Label Trial. Am J Psychiatry. 2025;182(1):104–13.

16. Erritzoe D, Barba T, Greenway KT, Murphy R, Martell J, Giribaldi B, et al. Effect of psilocybin versus escitalopram on depression symptom severity in patients with moderate-to-severe major depressive disorder: observational 6-month follow-up of a phase 2, double-blind, randomised, controlled trial. eClinicalMedicine. 2024;76.

17. Goodwin GM, Aaronson ST, Alvarez O, Arden PC, Baker A, Bennett JC, et al. Single-Dose Psilocybin for a Treatment-Resistant Episode of Major Depression. N Engl J Med. 2022;387(18):1637–48.

18. Falchi-Carvalho M, Palhano-Fontes F, Wießner I, Barros H, Bolcont R, Laborde S, et al. Rapid and sustained antidepressant effects of vaporized N,N-dimethyltryptamine: a phase 2a clinical trial in treatment-resistant depression. Neuropsychopharmacology: official publication of the American College of Neuropsychopharmacology. 2025;50(6):895-903.

19. Palhano-Fontes F, Barreto D, Onias H, Andrade KC, Novaes MM, Pessoa JA, et al. Rapid antidepressant effects of the psychedelic ayahuasca in treatment-resistant depression: a randomized placebo-controlled trial. Psychological Medicine. 2019;49(4):655–63.

20. 5-MeO-DMT for TRD clinical trial phase 2a 2024 [Available from: https://www.globenewswire.com/news-release/2024/03/27/2853041/0/en/atai-Life-Sciences-Announces-Positive-Initial-Results-from-Beckley-Psytech-s-Phase-2a-Open-Label-Study-of-BPL-003-Intranasal-5-MeO-DMT-in-Treatment-Resistant-Depression.html.

21. 5-MeO-DMT clinical trials phase 2b 2025 [Available from: https://investor.ghres.com/news-releases/news-release-details/gh-research-announces-primary-endpoint-met-phase-2b-trial-gh001.

22. COMP360 for PTSD clinical trial phase 2 2024 [Available from: https://ir.compasspathways.com/News--Events-/news/news-details/2024/Compass-Pathways-announces-durable-improvement-in-symptoms-through-12-weeks-in-open-label-phase-2-study-of-COMP360-psilocybin-in-post-traumatic-stress-disorder/default.aspx.

23. Ragnhildstveit A, Khan R, Seli P, Bass LC, August RJ, Kaiyo M, et al. 5-MeO-DMT for post-traumatic stress disorder: a real-world longitudinal case study. Frontiers in Psychiatry. 2023;Volume 14 - 2023.

24. Mitchell JM, Ot’alora G M, van der Kolk B, Shannon S, Bogenschutz M, Gelfand Y, et al. MDMA-assisted therapy for moderate to severe PTSD: a randomized, placebo-controlled phase 3 trial. Nature Medicine. 2023;29(10):2473–80.

25. Sessa B, Higbed L, O’Brien S, Durant C, Sakal C, Titheradge D, et al. First study of safety and tolerability of 3,4-methylenedioxymethamphetamine-assisted psychotherapy in patients with alcohol use disorder. Journal of Psychopharmacology. 2021;35(4):375–83.

26. Dodd S, Norman TR, Eyre HA, Stahl SM, Phillips A, Carvalho AF, et al. Psilocybin in neuropsychiatry: a review of its pharmacology, safety, and efficacy. CNS spectrums. 2023;28(4):416–26.

27. Hatzipantelis CJ, Olson DE. The Effects of Psychedelics on Neuronal Physiology. Annual Review of Physiology. 2024;86(Volume 86, 2024):27–47.

28. Ly C, Greb AC, Cameron LP, Wong JM, Barragan EV, Wilson PC, et al. Psychedelics Promote Structural and Functional Neural Plasticity. Cell Reports. 2018;23(11):3170–82.

29. Shao L-X, Liao C, Gregg I, Davoudian PA, Savalia NK, Delagarza K, et al. Psilocybin induces rapid and persistent growth of dendritic spines in frontal cortex in vivo. Neuron. 2021;109(16):2535–44.e4.

30. Gattuso JJ, Perkins D, Ruffell S, Lawrence AJ, Hoyer D, Jacobson LH, et al. Default Mode Network Modulation by Psychedelics: A Systematic Review. International Journal of Neuropsychopharmacology. 2022;26(3):155–88.

31. Siegel JS, Subramanian S, Perry D, Kay BP, Gordon EM, Laumann TO, et al. Psilocybin desynchronizes the human brain. Nature. 2024;632(8023):131–8.

32. Avram M, Fortea L, Wollner L, Coenen R, Korda A, Rogg H, et al. Large-scale brain connectivity changes following the administration of lysergic acid diethylamide, d-amphetamine, and 3,4-methylenedioxyamphetamine. Molecular psychiatry. 2025;30(4):1297–307.

33. Molendijk ML, de Kloet ER. Immobility in the forced swim test is adaptive and does not reflect depression. Psychoneuroendocrinology. 2015;62:389–91.

34. Hinchcliffe JK, Robinson ESJ. The Affective Bias Test and Reward Learning Assay: Neuropsychological Models for Depression Research and Investigating Antidepressant Treatments in Rodents. Current Protocols. 2024;4(6):e1057.

35. Stuart SA, Butler P, Munafo MR, Nutt DJ, Robinson ES. A translational rodent assay of affective biases in depression and antidepressant therapy. Neuropsychopharmacology: official publication of the American College of Neuropsychopharmacology. 2013;38(9):1625–35.

36. Harmer CJ, Duman RS, Cowen PJ. How do antidepressants work? New perspectives for refining future treatment approaches. The lancet Psychiatry. 2017;4(5):409–18.

37. Pringle A, Browning M, Cowen PJ, Harmer CJ. A cognitive neuropsychological model of antidepressant drug action. Progress in neuro-psychopharmacology & biological psychiatry. 2011;35(7):1586–92.

38. Stuart SA, Butler P, Munafo MR, Nutt DJ, Robinson ESJ. Distinct Neuropsychological Mechanisms May Explain Delayed-Versus Rapid-Onset Antidepressant Efficacy. Neuropsychopharmacology: official publication of the American College of Neuropsychopharmacology. 2015;40(9):2165–74.

39. Hinchcliffe JK, Stuart SA, Mendl M, Robinson ESJ. Further validation of the affective bias test for predicting antidepressant and pro-depressant risk: effects of pharmacological and social manipulations in male and female rats. Psychopharmacology (Berl). 2017;234(20):3105–16.

40. Stuart SA, Wood CM, Robinson ESJ. Using the affective bias test to predict drug-induced negative affect: implications for drug safety. British journal of pharmacology. 2017;174(19):3200–10.

41. The 3Hs Initiative [Available from: https://www.3hs-initiative.co.uk/.

42. Stuart SA, Robinson ES. Reducing the stress of drug administration: implications for the 3Rs. Sci Rep. 2015;5:14288.

43. Godlewska BR, Harmer CJ. Cognitive neuropsychological theory of antidepressant action: a modern-day approach to depression and its treatment. Psychopharmacology (Berl). 2021;238(5):1265–78.

44. Harmer CJ, Goodwin GM, Cowen PJ. Why do antidepressants take so long to work? A cognitive neuropsychological model of antidepressant drug action. The British Journal of Psychiatry. 2009;195:102–8.

45. Harmer CJ, O’Sullivan U, Favaron E, Massey-Chase R, Ayres R, Reinecke A, et al. Effect of acute antidepressant administration on negative affective bias in depressed patients. Am J Psychiatry. 2009;166(10):1178–84.

46. Barrett FS, Doss MK, Sepeda ND, Pekar JJ, Griffiths RR. Emotions and brain function are altered up to one month after a single high dose of psilocybin. Scientific Reports. 2020;10(1):2214.

47. Kometer M, Schmidt A, Bachmann R, Studerus E, Seifritz E, Vollenweider FX. Psilocybin Biases Facial Recognition, Goal-Directed Behavior, and Mood State Toward Positive Relative to Negative Emotions Through Different Serotonergic Subreceptors. Biological psychiatry. 2012;72(11):898–906.

48. de Vos CMH, Mason NL, Kuypers KPC. Psychedelics and Neuroplasticity: A Systematic Review Unraveling the Biological Underpinnings of Psychedelics. Frontiers in Psychiatry. 2021;Volume 12 - 2021.

49. Dolder PC, Schmid Y, Müller F, Borgwardt S, Liechti ME. LSD Acutely Impairs Fear Recognition and Enhances Emotional Empathy and Sociality. Neuropsychopharmacology: official publication of the American College of Neuropsychopharmacology. 2016;41(11):2638–46.

50. Wardle MC, de Wit H. MDMA alters emotional processing and facilitates positive social interaction. Psychopharmacology (Berl). 2014;231(21):4219–29.

51. Ramos L, Vicente SG. The effects of psilocybin on cognition and emotional processing in healthy adults and adults with depression: a systematic literature review. Journal of clinical and experimental neuropsychology. 2024;46(5):393–421.

52. Hinchcliffe JK, Kamenish K, Bartlett JM, Arban R, Hengerer B, Robinson ESJ. Modulation of negative affective biases in male rats may predict the rapid and sustained antidepressant effects of different NMDA antagonists. bioRxiv. 2025:2025.02.26.640447.

53. Kiilerich KF, Lorenz J, Scharff MB, Speth N, Brandt TG, Czurylo J, et al. Repeated low doses of psilocybin increase resilience to stress, lower compulsive actions, and strengthen cortical connections to the paraventricular thalamic nucleus in rats. Molecular psychiatry. 2023;28(9):3829–41.

54. Lima da Cruz RV, Costa RBGdM, de Queiroz GM, Stojanovic T, Moulin TC, Leão RN. Single-dose DMT reverses anhedonia and cognitive deficits via restoration of neurogenesis in a stress-induced depression model. Translational Psychiatry. 2026.

55. Brouns EJ, Ekins TG, Ahmed OJ. Single-dose psychedelic enhances cognitive flexibility and reversal learning in mice weeks after administration. Psychedelics. 2025;1(3):29–35.

56. Torrado Pacheco A, Olson RJ, Garza G, Moghaddam B. Acute psilocybin enhances cognitive flexibility in rats. Neuropsychopharmacology: official publication of the American College of Neuropsychopharmacology. 2023;48(7):1011–20.

57. Cameron LP, Olson DE. Dark Classics in Chemical Neuroscience: N,N-Dimethyltryptamine (DMT). ACS chemical neuroscience. 2018;9(10):2344–57.

58. Otto ME, van der Heijden KV, Schoones JW, van Esdonk MJ, Borghans LGJM, Jacobs GE, et al. Clinical Pharmacokinetics of Psilocin After Psilocybin Administration: A Systematic Review and Post-Hoc Analysis. Clinical Pharmacokinetics. 2025;64(1):53–66.

59. Reckweg JT, Uthaug MV, Szabo A, Davis AK, Lancelotta R, Mason NL, et al. The clinical pharmacology and potential therapeutic applications of 5-methoxy-N,N-dimethyltryptamine (5-MeO-DMT). J Neurochem. 2022;162(1):128–46.

60. Nelson DL, Lucaites VL, Audia JE, Nissen JS, Wainscott DB. Species differences in the pharmacology of the 5-hydroxytryptamine2 receptor: structurally specific differentiation by ergolines and tryptamines. The Journal of pharmacology and experimental therapeutics. 1993;265(3):1272–9.

61. Ling S, Ceban F, Lui LMW, Lee Y, Teopiz KM, Rodrigues NB, et al. Molecular Mechanisms of Psilocybin and Implications for the Treatment of Depression. CNS drugs. 2022;36(1):17–30.

